# Deep learning enables accurate soft tissue deformation estimation *in vivo*

**DOI:** 10.1101/2023.09.04.556266

**Authors:** Reece D. Huff, Frederick Houghton, Conner C. Earl, Elnaz Ghajar-Rahimi, Ishan Dogra, Denny Yu, Carisa Harris-Adamson, Craig J. Goergen, Grace D. O’Connell

## Abstract

Image-based deformation estimation is an important tool used in a variety of engineering problems, including crack propagation, fracture, and fatigue failure. These tools have been instrumental in biomechanics research where measuring *in vitro* and *in vivo* tissue deformations help evaluate tissue health and disease progression. However, accurately measuring tissue deformation *in vivo* is particularly challenging due to limited image signal-to-noise ratio. Therefore, we created a novel deep-learning approach for measuring deformation from a sequence of *in vivo* images called `StrainNet`. Utilizing a training dataset that incorporates image artifacts, `StrainNet` was designed to maximize performance in challenging *in vivo* settings. Artificially generated image sequences of human flexor tendons undergoing known deformations were used to compare `StrainNet` against two conventional image-based strain measurement techniques. `StrainNet` outperformed the traditional techniques by nearly 90%. High-frequency ultrasound imaging was then used to acquire images of the flexor tendons engaged during contraction. Only `StrainNet` was able to track tissue deformations under the *in vivo* test conditions. Findings revealed strong correlations between tendon deformation and contraction effort, highlighting the potential for `StrainNet` to be a valuable tool for assessing preventative care, rehabilitation strategies, or disease progression. Additionally, by using real-world data to train our model, `StrainNet` was able to generalize and reveal important relationships between the effort exerted by the participant and tendon mechanics. Overall, `StrainNet` demonstrated the effectiveness of using deep learning for image-based strain analysis *in vivo*.

## 1 Introduction

Image-based deformation measurement has been utilized in many engineering problems, such as crack propagation^1^, fracture^2^, and fatigue^3^. When applied to medical images, these techniques have aided in disease diagnostics^4–7^, assessment of injury mechanisms^8–11^, and evaluation of disease pathology^6,12–14^. Interest in using non-invasive approaches for tracking deformation *in vivo* has grown, due to its potential in designing safe work or appropriate rehabilitation strategies^15,16^. However, using medical images to accurately measure strain is difficult, partly due to image noise and limited resolution, where image artifacts can negatively impact accuracy^17–29^. Therefore, there is increasing importance for developing techniques that can perform well under clinical settings^9,30^.

Recent developments in machine learning have shown promise in measuring strains and this approach may present benefits over traditional strain measurement methods^31–34^. In particular, deep learning techniques, such as convolutional neural networks (CNNs), have been applied to predict strain maps between successive images. These methods resulted in more accurate and robust measurements compared to traditional image texture correlation techniques, such as digital image correlation (DIC), in controlled *in vitro* settings^32^. The training required for deep learning approaches also provides an advantage over traditional methods, because training the model with a large dataset of known strains allows the model to ignore image artifacts that can reduce accuracy^31–34^. Thus, the technique can be applied to a wider range of images, including those with lower signal-to-noise ratios, which can be challenging for traditional image texture correlation methods (**Section S1** in the Supplementary Information). Overall, the use of deep learning for image-based strain measurement has the potential to greatly improve assessment and understanding of *in vivo* tissue mechanics.

Here, we introduce a novel deep-learning approach, called `StrainNet`, specifically designed to maximize performance in challenging, *in vivo* settings. `StrainNet` utilizes a two-stage CNN architecture to predict full-field strain maps from a sequence of images that may be acquired in a medical setting (*e*.*g*., ultrasound images). The network was trained using a customized dataset, based on observations of tissue deformation and image artifacts *in vivo*, allowing it to overcome image artifacts that hamper traditional methods, such as image noise and artifacts, and provide accurate, full-field deformation predictions. We test and validate `StrainNet` on synthetic images with known deformations and real, experimentally collected ultrasound images of the flexor tendon in tension. Our results demonstrate that `StrainNet` outperforms traditional image texture correlation algorithms in both synthetic and real *in vivo* datasets, ultimately revealing novel strong correlations between tendon strains and applied forces as well as between load magnitude and measured mechanical properties *in vivo*. Our models and a tutorial for utilizing `StrainNet` are freely available at strainnet.github.io.

## 2Results

### 2.1 Human flexor tendons undergoing contraction

To evaluate `StrainNet’s` ability to predict tissue deformation, we designed a novel test configuration to measure grip forces in participants squeezing a dynamometer while simultaneously collecting *in vivo* images of their *flexor digitorum superficialis* (FDS) tendon (**Figure 1a.**; MicroFET, Hoggan Scientific, Salt Lake City, UT, United States). To begin, a participant’s forearm was secured to a custom platform that supported and stabilized the forearm and wrist allowing for hand exertions while a high-frequency ultrasound probe continuously collected images in the long axis throughout the protocol (**Figure 1a.**; Vevo3100 Ultrasound Imaging System, FUJIFILM VisualSonics Inc., Toronto, Ontario Canada; 21 MHz center frequency linear array ultrasound transducer; 15-30 MHz bandwidth; MX250). The participant was then asked to grip the dynamometer to their maximal effort to determine their maximum voluntary contraction (MVC). Averaged over the three trials, the participant’s MVC was 289.8N*±*19.8N. The participant was then asked to contract their forearm to three different effort levels—10%, 30%, and 50% of their MVC—by ramping up to the desired effort level over three seconds, holding the contraction for five seconds, then releasing the exertion to a full three seconds (**Figure 1a.**). Each effort level was repeated five times for a total of fifteen trials (*n*=15); however, data from two trials were lost due to corruption of the data file. All of the trials were performed with Purdue Institutional Review Board approval (IRB-2020-497).

**Figure 1.**
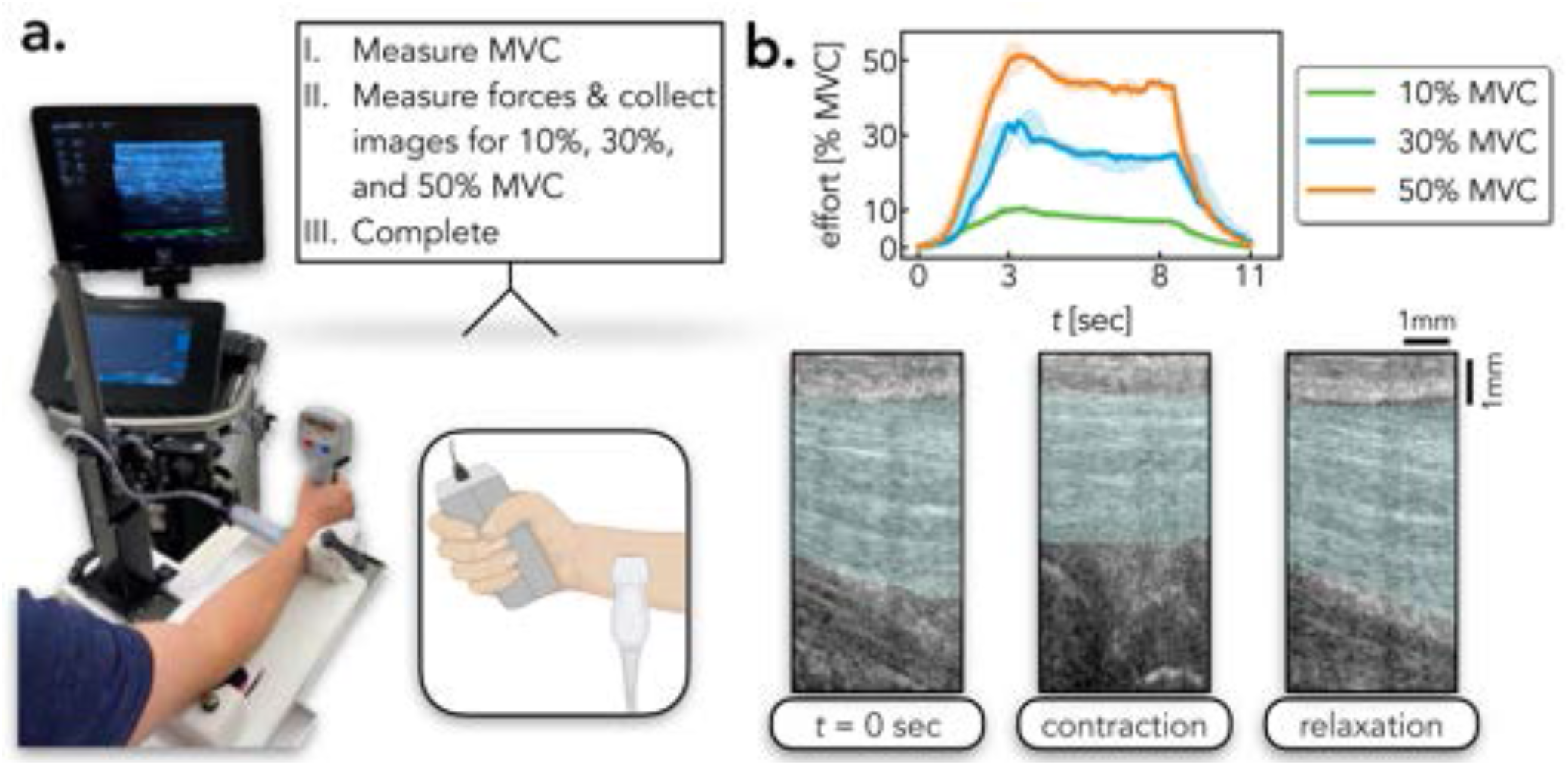
Experimental protocol, measured forces, and ultrasound images. **a**. Custom mount with the participant gripping the dynamometer to measure forces while high-frequency ultrasound images were collected. **b**. Once the participant’s maximum voluntary contraction (MVC) was determined, the participant was asked to squeeze the dynamometer to 10%, 30%, and 50% MVC. Data is shown throughout the contraction, hold, and relaxation for 10%, 30%, and 50% MVC. Characteristic images of the *flexor digitorum superficialis* (FDS) tendon, represented in teal, during the initiation of the test, during contraction (*i*.*e*., hold period), and after relaxation.

The participants maintained the target MVC throughout the 5 second hold period with only a maximum 16% difference between the desired and measured MVC (**Figure 1b.**; **Section S2** in the Supplementary Information). Over the course of the loading period, the tendon elongated and translated superficially before returning to its original position after relaxation (**Figure 1b.**). The testing configuration enabled robust evaluation of `StrainNet’s` performance in predicting tissue deformation under a range of physiological strains, including those that may be encountered during daily activities.

### 2.2 StrainNet outperforms traditional techniques in controlled environments

To test the accuracy of our strain analysis method in a controlled environment, five synthetic test cases were created by artificially imposing a non-linear strain field onto ultrasound images of the FDS tendon. The test cases simulated the experimental procedure with contraction, hold, and relaxation periods, as described above. The prescribed non-linear strain field was designed to reflect reported observations of *in vivo* tendon mechanics. Specifically, the strain in the superficial layer of the tendon was set to 75% of that in the deep layer^35^, and the tendon was modeled as an incompressible material^36,37^. The five test cases differed in their maximum longitudinal strain, 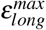 which was set to 4%, 7%, 10%, 13%, and 16% to cover the range of reported *in vivo* strains^35,37–39^. Noise representative of that present in ultrasound imaging was added to all synthetic test cases to emulate challenges present in the experimental dataset. A complete description of the synthetic test cases is provided in **Section S5**. By using synthetic test cases with known deformations, we were able to compare the performance and accuracy of our deep learning based approach with existing texture correlation algorithms.

`StrainNet` was benchmarked against two image-based strain algorithms, digital image correlation (DIC)^40^ and direct deformation estimation (DDE)^17^, using the synthetic test cases. `StrainNet` significantly outperformed the traditional texture correlation algorithms in all synthetic test cases; the median strain error from `StrainNet` was 48-84% lower than the strain error from both DIC and DDE (**Figure 2a.**; *p*<0.001 in all strain cases). In addition to the overall performance comparison, temporal analysis of strain error further highlights the advantages of `StrainNet` (**Figure 2b.**). The accuracy of `StrainNet` was nearly 90% better than DIC and DDE across all test cases (solid lines in **Figure 2b.**). `StrainNet` was also 90% more precise than DDE; however, DIC was the most precise algorithm tested (filled-in area in **Figure 2b.**).

**Figure 2.**
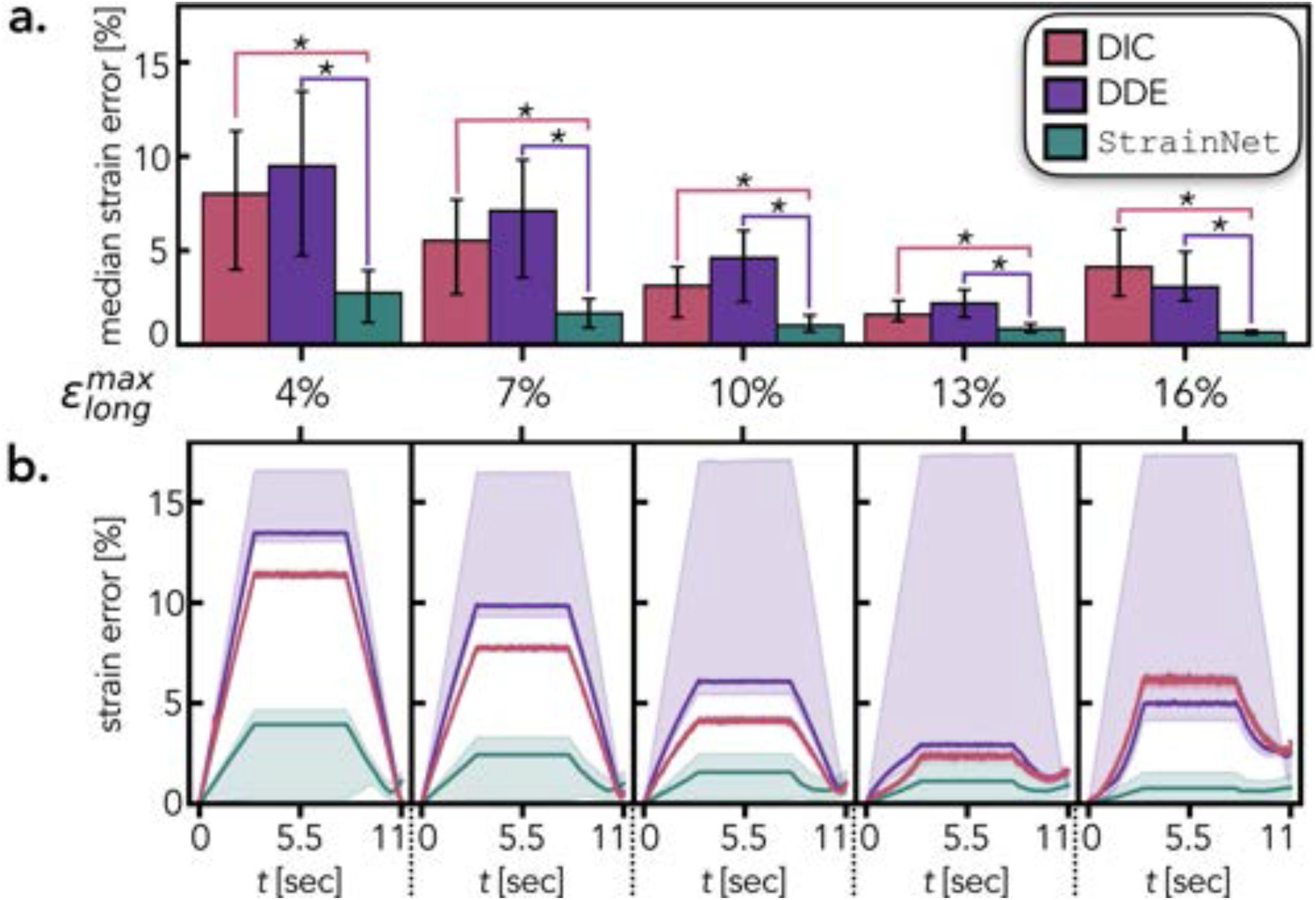
Quantitative evaluation of performance of DIC, DDE, and StrainNet on synthetic test cases. **a**. Median strain error calculated over all ultrasound frames. Error bars indicate the first and third quartiles of the strain error. The asterisks denote a statistically significant difference between connected groups (*p* < 0.001). **b**. Temporal strain error for each synthetic test case. The solid line indicates the median strain error for each ultrasound frame and the shadded area shows the first and third quartile range of the spatial strain error.

`StrainNet` achieved pixel-wise strain estimation, while DIC and DDE were limited to the central area of interest (**Figure 3a.**). DDE and `StrainNet` were able to accurately capture the heterogeneous nature of the applied strain, whereas the DIC-predicted strain field was homogeneous (**Figure 3a.**). The DIC analysis area was limited to within the boundaries of the tendon whereas DDE and `StrainNet` cover both the tendon and the surrounding soft tissue, revealing large (10%) spatial strain error at the boundary (**Figure 3b.**). All three methods exhibited low spatial strain error throughout the tendon during contraction and relaxation (**Figure 3b.**).

**Figure 3.**
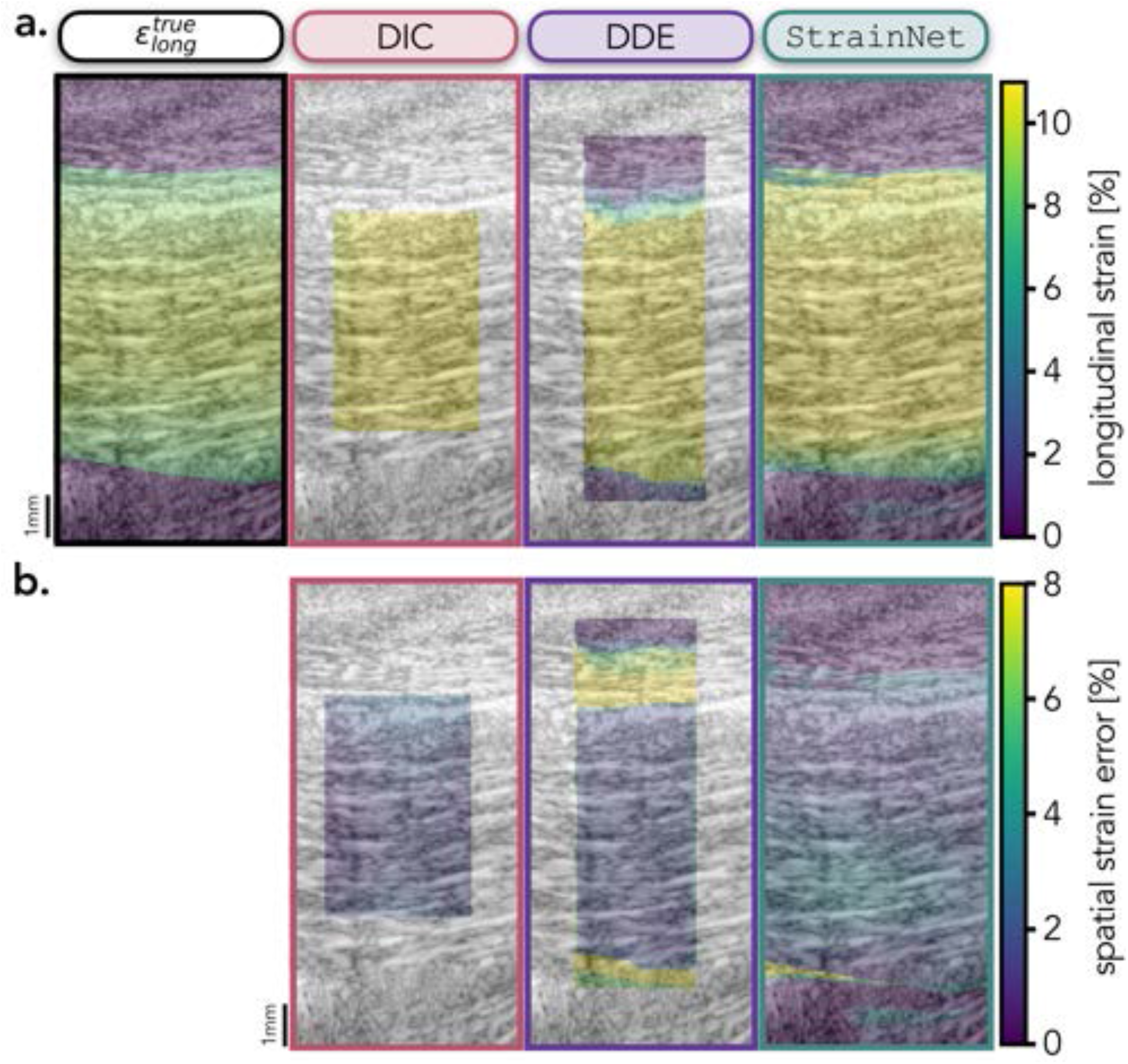
Qualitative evaluation of DIC, DDE, and StrainNet’s performance on the synthetic test case (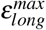 = 10%). **a**. From left to right: the true prescribed longitudinal strain followed by DIC-, DDE-, and `StrainNet`-predicted strain fields during the hold period. **b**. From left to right: spatial distributions of strain error for DIC-, DDE-, and `StrainNet` during the hold period.

### 2.3 StrainNet enables accurate *in vivo* deformation estimation

Both DIC and DDE had difficulties tracking tissue deformations from *in vivo* images and many pixels were lost during analysis. `StrainNet`, on the other hand, was able to learn around much of the noise and predict the longitudinal strain in the tendon, which increased with the effort exerted by the participant (**Figure 4a.**). There was a strong linear relationship between the `StrainNet`-predicted longitudinal strain and effort level (**Figure 4b.**; *r* = 0.78, *p* = 0.002). Similarly, there was also a strong linear relationship between the apparent modulus calculated with the `StrainNet`-predicted strain and effort level (**Figure 4c.**; *r* = 0.87, *p* < 0.001).

**Figure 4.**
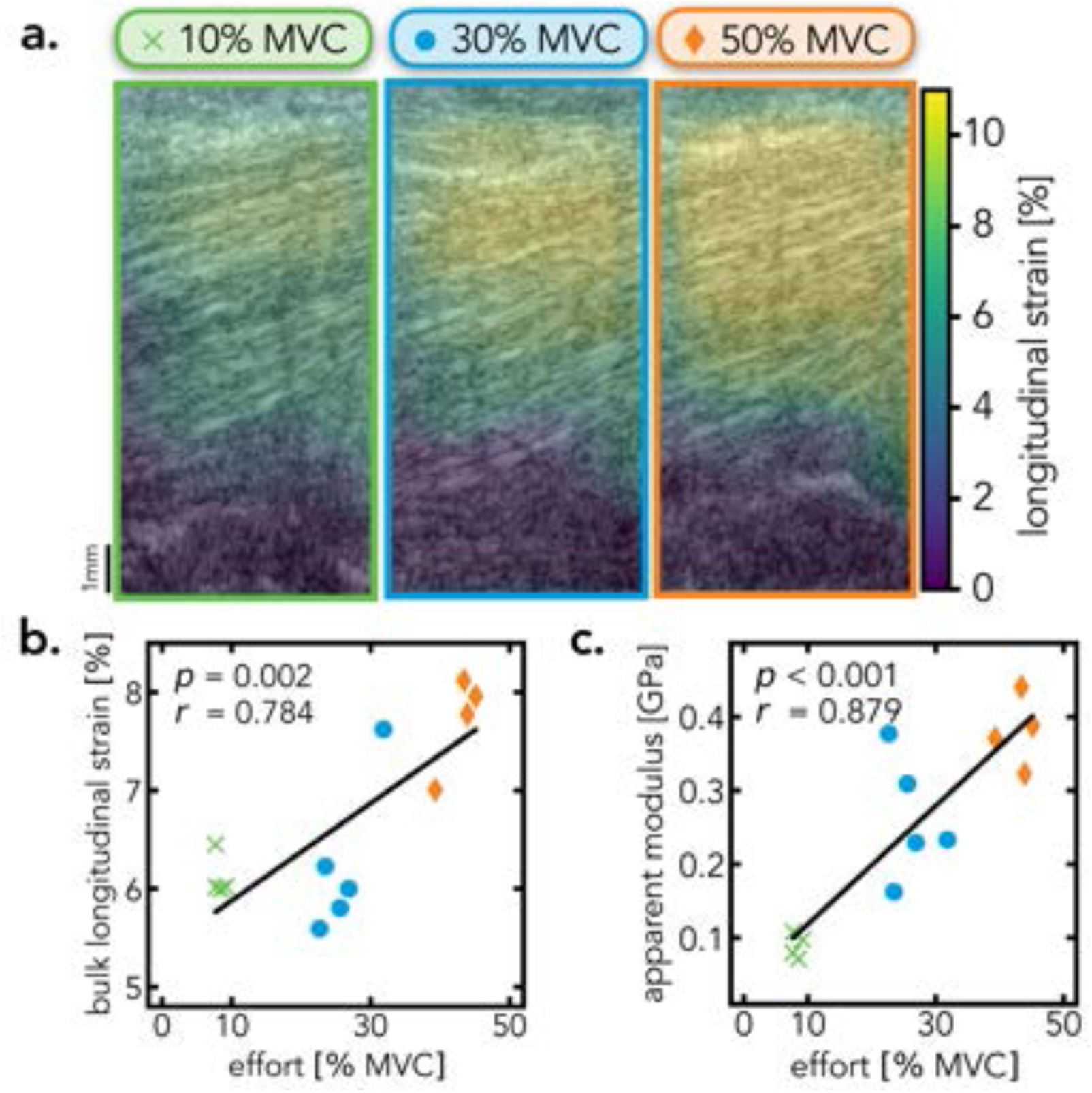
Quantitative and qualitative analysis of StrainNet applied to *in vivo* images. **a**. `StrainNet`-measured spatial distribution of longitudinal strain throughout the tendon during the contraction to 10%, 30%, and 50% MVC. **b**. Linear regression between the bulk longitudinal strain in the tendon and the effort exerted by the participant. **c**. Linear regression between apparent modulus and effort level.

## 3 Discussion

`StrainNet` was able to accurately measure different strain levels using ultrasound images of the flexor tendon. For synthetic datasets, `StrainNet` detected subtle differences in deformations with a high degree of accuracy (< 3% error), outperforming existing approaches (*e*.*g*., DIC and DDE) that had median strain errors as high as 10%. Additionally, when applied to *in vivo* ultrasound images, `StrainNet` predicted a strong linear correlation between the measured strain and effort level (percentage of the MVC), further validating the performance of the model. Finally, full-field deformation predictions were able to unveil stress-strain curves, and thus measure mechanical properties within soft biological tissue under physiological boundary conditions. Taken together, these findings suggest that deep learning models have the potential to significantly advance the accuracy of *in vivo* biomechanics studies.

There are several limitations to our model that will be addressed in future work. First, the model was evaluated on a single tissue type and location, so it is not clear whether it can be applied to a wider range of tissue types. Additionally, the current architecture is specialized to handle only three types of deformation, and it would be useful to explore expanding its capabilities to a wider range of deformations (*e*.*g*., shear). However, given the customizability of the training set (**Section S4** in the Supplemental Information) and the robustness of `StrainNet`, model alignment may lead to accurate measurement of deformation in other biological tissues.

The potential applications of `StrainNet` are vast and promising. `StrainNet` significantly surpassed traditional image texture correlation methods in controlled environments, such as synthetic test cases (**Figure 2**). Moreover, in more complex settings where image texture correlation is susceptible to errors caused by image artifacts, `StrainNet` consistently delivered accurate and expected tissue deformation levels (**Figure 4**), in line with previous reports^38,39,41^. Furthermore, the measured tissue mechanical properties aligned with those previously reported for human patellar and Achilles tendons with similar experimental procedures *in vivo* (**Section S6** in the Supplemental Information)^41–45^. Taken together, our results suggest that `StrainNet` can be employed in a broad array of biomedical applications, such as *in vivo* imaging studies of muscle function, blood flow, and tissue viability. In summary, the design and capabilities of `StrainNet` hold immense potential for quantifying biomechanical metrics, leading to substantial progress in assessing soft tissue deformation.

## 4 Methods

### 4.1 StrainNet architecture & training

The `StrainNet` architecture was specifically designed to handle the unique challenges presented with *in vivo* image analysis. Specifically, `StrainNet` was developed and trained to predict strain within high-frequency ultrasound images of FDS tendons undergoing contraction, as described above (**Section 2.1**). The architecture was constructed to first classify the image pair as undergoing tension, compression, or rigid body motion, and then to apply an appropriate neural network to predict the strain field within the tendon (**Figure 5a.**).

**Figure 5.**
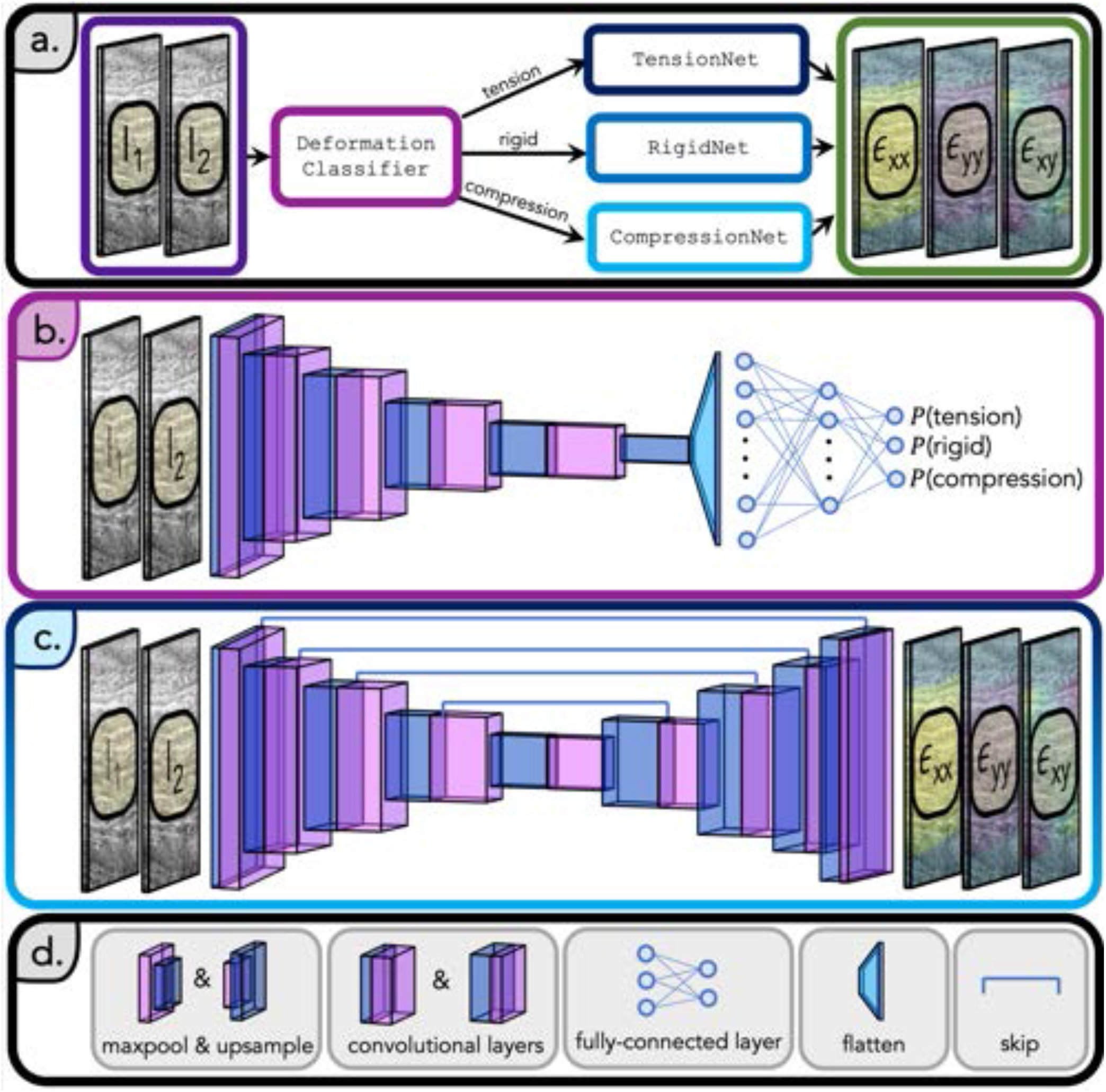
Architecture of StrainNet. **a**. `StrainNet` includes a deep neural network trained to predict a relationship between input images, *I*_1_ and *I*_2_ and tendon strain, *ε*_*xx*_, *ε*_*yy*_, *ε*_*xy*_. `StrainNet` comprises two stages, where the first stage was the `DeformationClassifier` and the second stage included `TensionNet`, `CompressionNet`, and `RigidNet`. **b**. `DeformationClassifier` is composed of convolutional layers, max pooling, and Rectified Linear Unit (ReLU) activation functions. The resulting features are flattened and passed through a fully-connected neural network to predict the probability of the image pair undergoing tension, compression, or rigid body motion. **c**. The architecture of `TensionNet`, `CompressionNet`, and `RigidNet` included convolutional layers, max pooling, upsampling, skip layers, and ReLU activation functions to predict the full strain field (*ε*_*xx*_, *ε*_*xy*_, *ε*_*yy*_). **d**. Blocks in **b**. and **c**. were connected by ReLU activation functions and utilized batch normalization.

The first stage of the architecture, the `DeformationClassifier`, was a CNN that classifies the image pair based on the type of deformation (**Figure 5a.**). It consisted of a series of convolutional layers, max-pooling layers, and fully connected layers. The convolutional layers extracted features from the image, while the max-pooling layers reduced dimensionality. The fully connected layers were used to make the final classification (**Figure 5b.**).

Once the image pair was classified, it was passed to one of three neural networks: `TensionNet`, `CompressionNet`, or `RigidNet` (**Figure 5a.**). These networks predicted the strain field from the input image pair and were based on the UNet architecture, a popular biomedical image segmentation deep-learning architecture^46^. The architecture utilized an encoder-decoder structure, with the encoder extracting features and the decoder up-sampling feature maps to the original image size. The encoder and decoder in `TensionNet`, `CompressionNet`, and `RigidNet` were composed of convolutional layers, max-pooling layers, up-sampling layers, and Rectified Linear Unit (ReLU) activation functions (**Figure 5c.-d.**). Skip connections between the encoder and decoder were included to help improve strain field prediction quality by reducing vanishing gradients^47^.

To effectively train `StrainNet`, a diverse training set was created with image pairs of deformation fields that emulated real-world observations and image artifacts commonly encountered in medical imaging (*e*.*g*., random noise). The training set included 3,750 synthetically generated image pairs and 1,250 experimental image pairs. Training set generation involved utilizing a generalized mathematical model of tendon mechanics, prescribing the non-linear strain fields onto collected ultrasound images of the tendon, and adding noise to simulate real-world imaging conditions. Deformation and noise parameters were randomly sampled from uniform probability distributions, ensuring a robust dataset for learning the strain measurement task. The detailed process of generating the training set, including the acquisition of experimental data, image preprocessing, and the combination of synthetic and experimental examples, can be found in **Section S4** in the Supplemental Information.

Following the creation of the training set, the `StrainNet` model was trained using a combination of loss functions tailored to the specific tasks of each subnetwork. For the `DeformationClassifier`, a cross-entropy loss function was utilized and defined as

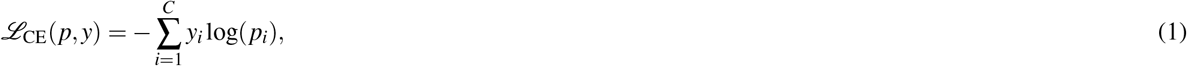

where *p* represents the predicted class probabilities, *y* is the true one-hot encoded class label, and *C* is the number of classes (tension, rigid, and compression). For the other three models, `TensionNet`, `CompressionNet`, and `RigidNet`, the mean *ℓ*_2_ loss function was used and expressed as

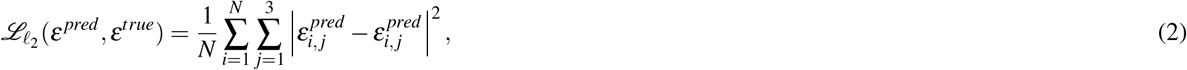

where 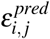 and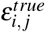 denote *i*-th sample and the *j*-th component (longitudinal, transverse, and shear across all image pixels) of the predicted and true strain field, respectively, and *N* is the number of examples in the training batch.

The training process was conducted for 100 epochs using the Adam optimizer (PyTorch^48^ 1.12.1) on an NVIDIA K100 16GB graphics processing unit (GPU) with a learning rate of 0.001. Different batch sizes were employed for the sub-models to accommodate their specific training requirements. For the `DeformationClassifier`, a batch size of 100 was used to take advantage of parallel processing and to reduce the noise in gradient updates. In contrast, a smaller batch size of 10 was utilized for the `TensionNet`, `CompressionNet`, and `RigidNet` models, allowing for more frequent weight updates and improved convergence properties. Training of the `DeformationClassifier` required 4 hours, whereas `TensionNet`, `CompressionNet`, and `RigidNet` needed approximately 8 hours to train. The combination of these hyperparameters, the GPU, and the optimizer facilitated successful training of `StrainNet`, enabling it to learn the relationships between ultrasound images of tendons and their corresponding strain fields.

### 4.2 Strain analysis method validation

`StrainNet’s` performance and accuracy was compared to two existing texture correlation algorithms, including DIC^40^ and DDE^17^. All three techniques were applied to all synthetic test cases. Given the applied straitensor, the spatial strain error was calculated as

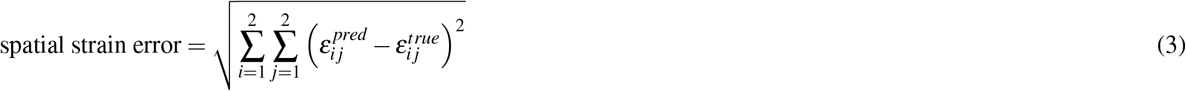

where 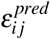 and 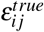 represent the true and predicted strain tensor for all pixels in each ultrasound image. To robustly evaluate the performance, strain error and median strain error were calculated as

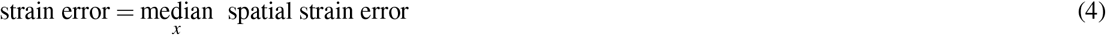

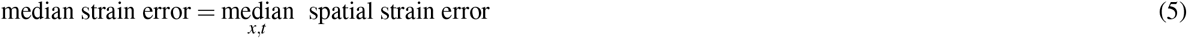

where the underset *x* and *x,t* denote that the median was obtained across the entire ultrasound image or all ultrasound images throughout the full contraction-relaxation cycle, respectively. To compare the strain error for each synthetic test case between `StrainNet` and DIC, as well as between `StrainNet` and DDE, permutation tests of the strain errors were conducted for each test case.

### 4.3 Experimental strain analysis and mechanical property estimation

`StrainNet`, DIC, and DDE were then applied to the experimental images. To quantify the bulk tendon mechanical behaviour, the bulk longitudinal strain during the hold period was calculated as the median longitudinal strain over the tendon region

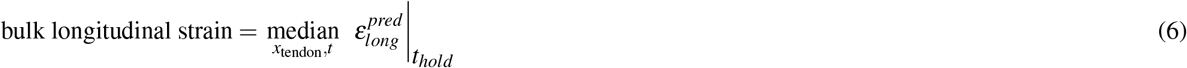

where *x*_tendon_ and *t*_*hold*_ represent the region of the image containing the tendon and the time over the hold period, respectively. Linear regression was performed to examine the relationship between the effort level and the corresponding bulk longitudinal strain for each of the three methods (`StrainNet`, DIC, and DDE).

The bulk longitudinal strains were then used to estimate the mechanical properties of the tendon. First, the longitudinal tendon stress was calculated by dividing the measured force by the tendon’s cross-sectional area, which was manually segmented from ultrasound images. Subsequently, the tendon’s apparent modulus was calculated as the slope of the linear region of each trial’s stress-strain curve. Finally, linear regression on the measured apparent modulus and the effort level was performed.

The significance level for all permutation and linear regression tests was set to 0.05.

## Data Availability

The pre-trained models are available on the project page, strainnet.github.io.

## Code Availability

The code is publicly available at github.com/reecehuff/StrainNet. The project page, strainnet.github.io, also includes a detailed tutorial for implementing `StrainNet` in any desired experimental setup with any biological tissue. All data and code questions and requests should be addressed to R.D.H. at.

## Acknowledgements

R.D.H. was funded by a fellowship from the National Science Foundation (NSF GRFP; DGE 2146752). This study was also supported by the National Institutes of Health (NIH R21 AR075127-02) and the National Institute for Occupational Safety and Health (NIOSH) & Centers for Disease Control and Prevention (CDC) (Training Grant T42OH008429). The authors also acknowledge Guoyang Zhou, Andrew J. Darling, and Frederick W. Damen for their valuable contributions to image analysis.

## Author contributions statement

R.D.H., F.H., D.Y., C.H.A., C.J.G., and G.D.O. conceptualized the study. C.C.E. and E.G.R. collected the experimental data. R.D.H., F.H., and I.D. processed the data. R.D.H. created the illustrations and figures. All authors aided in data interpretation. R.D.H. drafted the manuscript, and F.H., C.H.A., C.J.G., and D.Y. critically revised it. All authors approved the final manuscript.

