## Supplementary Information for "Deep learning enables accurate soft tissue deformation estimation *in vivo*"

#### Article

bioRxiv
THE PREPRINT SERVER FOR BIOLOGY

Section S5 Synthetic test cases 10

Section S6 Reported *in vivo* tendon modulus 12

References 15

### S1 Overview of image texture correlation and `StrainNet`

#### S1.1 Image texture correlation

Image texture correlation—or optical flow—is a prevalent image processing technique, originating in the 1980s [1–3]. This method fundamentally hinges on determining the displacement field between pixels of a reference (or "undeformed") image and a "deformed" image (Figure S1a.). This is achieved by employing a cost function, which can take the form of either a correlation or a loss function. The technique presupposes that pixel values, such as intensity, remain constant between the undeformed and deformed images. Two prominent variants of image texture correlation are described below.

##### S1.1.1 Digital Image Correlation (DIC)

Digital image correlation (DIC) is an extensively utilized image texture correlation method. It operates by dividing the reference images into a grid of rectangular boxes. The size of the box is often denoted as the subset size, while the spacing between the subsets is commonly referred to as the step size (Figure S1b.). DIC employs the cost function

$$\mathcal{C}_{\mathrm{DIC}}[\mathbf{U}_i] = \int_{\Omega_i} \left[\mathrm{I}_1(\mathbf{X}_i) - \mathrm{I}_2(\mathbf{X}_i + \mathbf{U}_i)\right]^2$$

where $\mathrm{I}_1$ and $\mathrm{I}_2$ are the reference and deformed images, respectively, and $\mathbf{X}_i$ and $\mathbf{u}_i$ represents the pixel coordinates and displacement centered in subset $\Omega_i$. Cost functions can be applied either locally (independently for each subset) or globally (enforcing displacement continuity among neighboring subsets). The discrete displacement field is then numerically differentiated to obtain a discrete strain field (Figure S1b.). Palanca *et al.* provides a comprehensive review of DIC applications in biomechanical engineering.

##### S1.1.2 Direct Deformation Estimation (DDE)

Direct deformation estimation (DDE) is similar to DIC in many aspects, but with a crucial difference: instead of calculating the displacement field, DDE determines the deformation gradient, $\mathbf{F}$, which maps the undeformed and deformed images via $\mathrm{d}\mathbf{x} = \mathbf{F}\mathrm{d}\mathbf{X}$ (Figure S1c.). The cost function utilized in DDE is of the form

$$\mathcal{C}_{\mathrm{DDE}}[\mathbf{U}_i] = \int_{\Omega_i} \left[\mathrm{I}_1(\mathrm{d}\mathbf{X}_i) - \mathrm{I}_2(\mathrm{d}\mathbf{x}_i + \mathbf{F}_i\mathrm{d}\mathbf{X}_i))\right]^2$$

where $\mathrm{d}\mathbf{X}_i$ and $\mathrm{d}\mathbf{x}_i$ denotes the change in pixel coordinates in the reference and deformed image, respectively, and $\mathbf{F}_i$ represents the deformation gradient in subset $\Omega_i$. By directly computing the strain from the deformation gradient without the need for displacement differentiation, DDE has been shown to provide better accuracy, noise insensitivity, and precision compared to displacement-based techniques like DIC [5].

##### S1.1.3 Limitations of image texture correlation

Despite the widespread use and success of image texture correlation techniques, such as digital image correlation (DIC) [6] and direct deformation estimation (DDE) [5, 7], they are not without their limitations. Some of the main drawbacks of these techniques include noise sensitivity, the need for sufficient texture, and the inability to handle large deformations. These limitations are discussed in detail below.

**Noise Sensitivity:** One of the significant limitations of image texture correlation techniques is their sensitivity to image noise [8]. The fundamental assumption behind these methods is that the image intensity remains constant between the undeformed and deformed images [9]. However, in real-world scenarios, images are often affected by various types of noise, such as sensor noise or environmental noise [10]. This noise can lead to errors in the estimation of displacement and strain fields, ultimately affecting the accuracy of the results [11, 12]. Several strategies have been proposed to mitigate the effect of noise, such as the use of pre-processing techniques to denoise images [13], incorporating regularization terms in the cost function [14], and employing robust optimization algorithms [15].

**Texture requirement:** Adequate image texture is a prerequisite for accurate displacement and deformation measurements using image texture correlation[16]. When images present low or uniform texture, the accuracy of these techniques can be compromised. To mitigate this, researchers have proposed the use of artificial speckle patterns or markers to increase image texture [17]. Also, the adoption of advanced image processing algorithms, capable of extracting subtle texture details, has been recommended [18]. Ensuring the overall accuracy of image correlation techniques under varied textures is pivotal for enhancing their applicability and precision across various fields [16–18].

**Large deformation handling:** Traditional image texture correlation techniques often struggle to handle large deformations or complex motion between the undeformed and deformed images. This limitation arises due to the linear approximation employed in the cost functions, which may not accurately capture the non-linear deformations present in the images [19]. To overcome this issue, researchers have developed multi-scale and multi-resolution approaches, where the deformation estimation is performed iteratively at various scales and resolutions, starting from a coarse scale and gradually refining the estimate at finer scales [20].

In conclusion, while image texture correlation techniques have proven to be valuable tools for deformation and displacement estimation, they have several limitations. Researchers have been continuously working to address these challenges and develop more advanced and robust techniques for a wide range of applications.

#### S1.2 Advances in machine learning

In recent years, the development of advanced algorithms and computational methods has greatly improved the performance of image-based deformation measurement techniques [21, 22]. Deep learning techniques have been introduced to enhance the pattern matching process, enabling more accurate and robust displacement estimations even in the presence of noise and other distortions [23]. Additionally, real-time processing capabilities are being developed to facilitate faster, more efficient measurements [24].

Recent developments in machine learning have shown promise in measuring strains from images over traditional strain measurement methods. In particular, deep learning techniques, such as convolutional neural networks (CNNs), have been applied to predict strain maps between successive images [25–27]. These methods have been shown to provide more accurate and robust results compared to traditional image texture correlation techniques, such as DIC, in controlled settings [25]. One of the key advantages of using deep learning for image-based strain measurement is that the model can be trained on a large dataset of image pairs with known strains, allowing the model to "learn" to overcome image artifacts that can negatively impact the accuracy of traditional techniques [25].

#### S1.3 `StrainNet`

Considering the limitations of image texture correlation and recent advancements in machine learning, we introduce a novel deep-learning approach called `StrainNet`, specifically designed to maximize performance in challenging, *in vivo* settings. `StrainNet` leverages the power of CNNs to accurately predict full-field strain maps from image pairs captured during *in vivo* imaging (**Figure S1d.**). Its robustness and adaptability allow it to overcome challenges such as low image quality and complex movements, making it a valuable tool for biomechanical research.

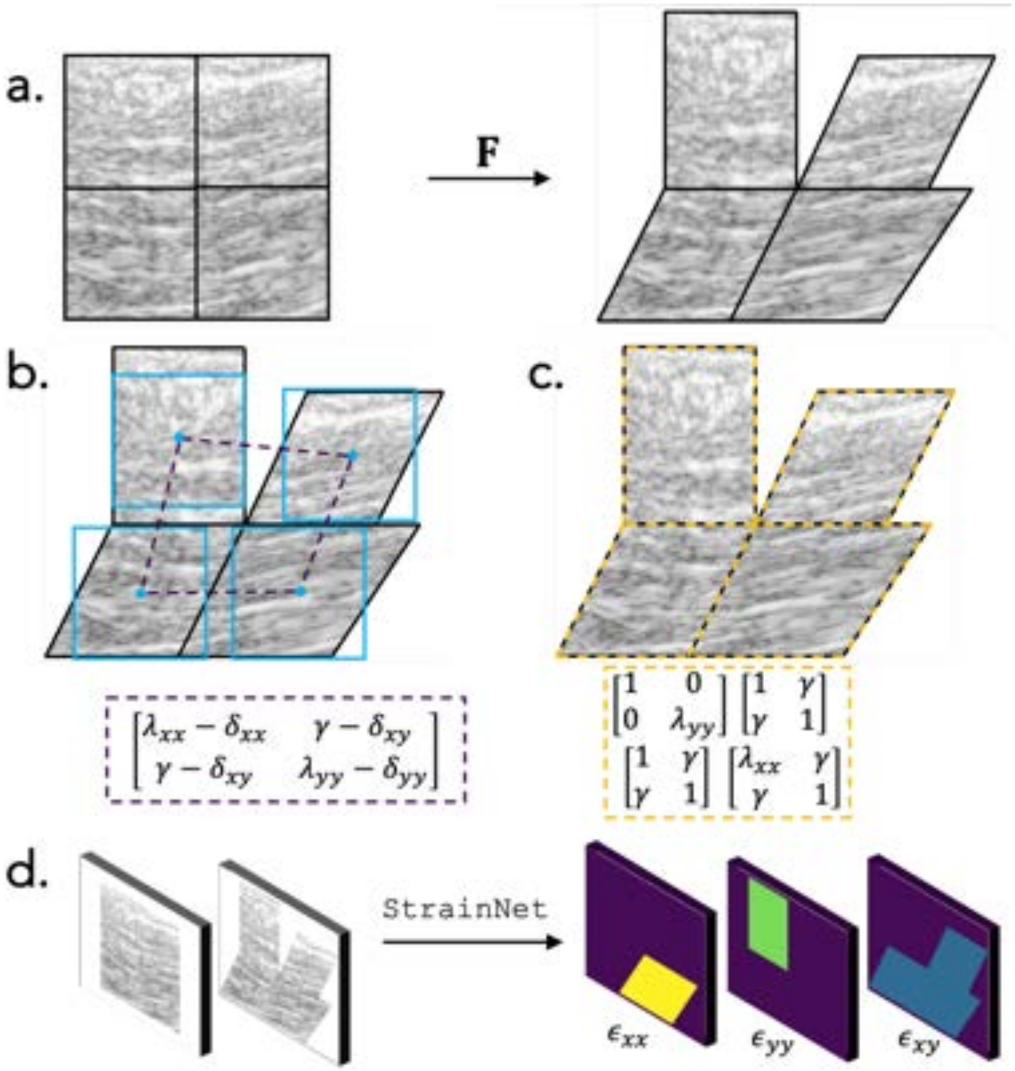

**Figure S1: Schematic illustrating the process of strain calculation using DIC, DDE, and `StrainNet` from an image pair. a.** The left image represents a reference image (i.e., $I_1$), and the right image represents a deformed image (i.e., $I_2$) exhibiting vertical tension in the top left ($\lambda_{yy}$), pure shear in the top right and lower left ($\gamma$), and a combination of shear with horizontal extension in the lower right corner ($\lambda_{xx}$ and $\gamma$). **b.** DIC solves for displacements of four pixels using square subset regions (blue boxes) and employs numerical differentiation to estimate strain (dark purple dashed box). $\delta_{\alpha\beta}$ represents the errors from numerical differentiation. **c.** DDE solves for the deformation gradient of each subset directly (orange dashed box). **d.** `StrainNet` estimates full-field strain given a pair of input images ($I_1$ and $I_2$).

#### S2 Measured error of effort level

To evaluate the discrepancy between the desired and measured effort levels during the experiments, we computed the percent root-mean-square error (RMSE) of the effort levels for 10%, 30%, and 50% of the participant's maximum voluntary (MVC) trials. This quantifies the deviation of the measured effort from the desired percentage of MVC. The percent error was defined as

$$\%\mathrm{RMSE} = \left[\frac{1}{3}\sum_{X\in[10,30,50]} \mathrm{RMSE}(X\%\ \mathrm{MVC})\right] \times 100$$

where

$$\mathrm{RMSE}(X\%\ \mathrm{MVC}) = \sqrt{\frac{1}{N_{X\%\ \mathrm{MVC}}} \sum_{i\in X\%\ \mathrm{MVC}} \left(\frac{y_i^{\mathrm{desired}} - y_i^{\mathrm{measured}}}{y_i^{\mathrm{desired}}}\right)^2}$$

where $y_i^{\mathrm{desired}}$ represents the desired effort level for the $i$-th trial at X% MVC, $y_i^{\mathrm{measured}}$ denotes the measured effort level for the same trial, and $N_{X\%\ \mathrm{MVC}}$ indicates the number of trials at X% MVC.

The results of this analysis are presented in Figure S2. The red bars depict the error between the measured effort and the desired percentage of MVC for each trial. The mean RMSE over all effort levels was found to be 16%, indicating that the measured effort levels were in close agreement with the requested MVC levels.

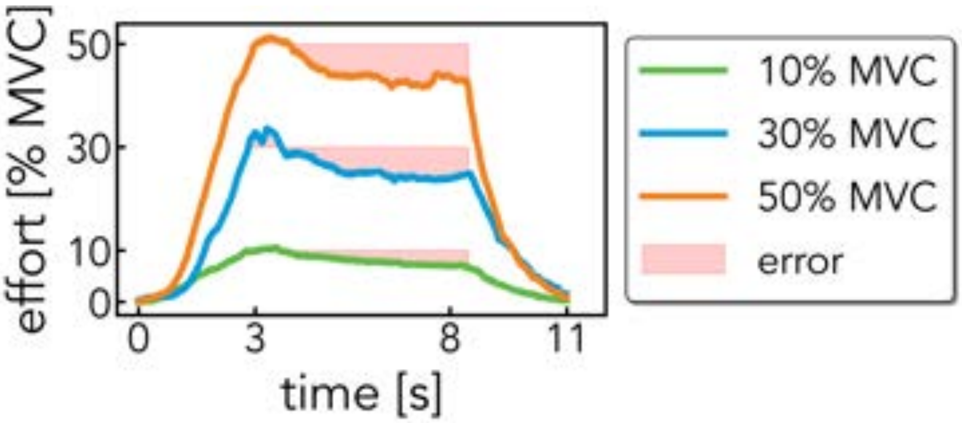

**Figure S2: Measured effort level in a participant average over trials.** The error between the measured effort and the desired percentage of the MVC is shown in red to facilitate visualization.

### S3 Tendon deformation model and image processing workflow

In order to train, benchmark, and tune `StrainNet`, an image-processing workflow was developed to induce known deformation within ultrasound images of tendons. This workflow was designed for broad applicability and to emulate observed *in vivo* tendon strain.

#### S3.1 Generalized mathematical model

Prior research has shown that deep tendon layers exhibit less strain than superficial layers [28]. Thus, a generalized strain field was specified along the tendon's length as

$$\epsilon_{long}^{max}(y) = \frac{4(\epsilon_{long}^{superficial} - \epsilon_{long}^{deep})(y - y_c)^2}{h_{tendon}^{BB}} + \epsilon_{long}^{deep} \tag{1}$$

where $h_{tendon}^{BB}$ denotes the tendon's width (**Figure S3a.**), $\epsilon_{long}^{superficial}$ represents the longitudinal strain applied at the superficial layer, and $\epsilon_{long}^{deep}$ signifies the longitudinal strain applied at the deep layer (**Figure S3b.**).

While the strain distribution in Equation (1) characterizes tendon spatial variation, the magnitude remains uncertain. Previous studies demonstrate that tendon longitudinal strain magnitudes range between 4% and 14% during contraction [28–31] under various loading conditions. However, both the load applied and the time allowed reach the load dictate the strain magnitudes within sequential images. Consequently, the time-dependence in the prescribed strain was expressed as

$$\epsilon_{long}^{true}(y, t) = \alpha t \epsilon_{long}^{max}(y, t) \tag{2}$$

where $t$ represents time in seconds and $\alpha$ is a parameter dictating the time taken for the deformation to attain its maximum. For instance, if a loading scenario reached the maximum strain in three seconds, then $\alpha$ would be set to $\frac{1}{3\text{seconds}}$ to account for time dependence.

To elucidate the relationships between the longitudinal strain $\epsilon_{long}^{true}$ defined in Equation (2) and the transverse strain $\epsilon_{trans}^{true}$, the shear strain $\epsilon_{shear}^{true}$, and the displacement field $\vec{u} = [u_x, u_y]$, a constant $k$ is defined as

$$k = \frac{4(\epsilon_{long}^{superficial} - \epsilon_{long}^{deep})}{h_{tendon}^{BB}}. \tag{3}$$

Hence, Equation (1) can be rewritten as

$$\epsilon_{long}^{max}(y) = k(y - y_c)^2 + \epsilon_{long}^{deep}. \tag{4}$$

By applying basic elasticity principles, the transverse strain field, $\epsilon_{trans}^{true}$, can be rewritten as

$$\epsilon_{trans}^{max}(y) = -\nu \epsilon_{long}^{max} = -\nu \left[k(y - y_c)^2 + \epsilon_{long}^{deep}\right] \tag{5}$$

The displacement field, $\vec{u} = (u_x, u_y)$, can be obtained by integrating Equation (4) and (5) with respect to $x$ and $y$, respectively.

$$u_x^{max}(x, y) = kx(y - y_c)^2 + \epsilon_{long}^{deep} x + C_1 = kx(y - y_c)^2 + \epsilon_{long}^{deep} x \tag{6}$$

$$u_y^{max}(y) = -\nu \left[\frac{k(y - y_c)^3}{3} + \epsilon_{long}^{deep} y\right] + C_2 = -\nu \left[\frac{k(y - y_c)^3}{3} + \epsilon_{long}^{deep}(y - y_c)\right] \tag{7}$$

It should be noted that $C_1 = 0$ is set such that the displacements are small near the distal end of the tendon where $x$ is small, and $C_2 = -\epsilon_{long}^{deep} y_c$ is set such that $u_y = 0$ when $y = y_c$. The shear strain in the tendon then becomes

$$\epsilon_{shear}^{max}(x, y) = \frac{1}{2}[u_{x,y} + u_{y,x}] = kx(y - y_c). \tag{8}$$

Therefore, the maximum applied strain field applied to the tendon is defined as

$$\epsilon_{long}^{max}(y) = k(y - y_c)^2 + \epsilon_{long}^{deep}, \tag{9}$$

$$\epsilon_{trans}^{max}(y) = -\nu \left[k(y - y_c)^2 + \epsilon_{long}^{deep}\right], \tag{10}$$

$$\epsilon_{shear}^{max}(x, y) = kx(y - y_c) \tag{11}$$

where $k = 4(\epsilon_{long}^{superficial} - \epsilon_{long}^{deep})/h_{tendon}^{BB}$. The corresponding maximum displacement field for this strain field is

$$u_x^{max}(x, y) = kx(y - y_c)^2 + \epsilon_{long}^{deep} x \tag{12}$$

$$u_y^{max}(y) = -\nu \left[\frac{k(y - y_c)^3}{3} + \epsilon_{long}^{deep}(y - y_c)\right] \tag{13}$$

The generalized time-dependent strain and displacement field can be expressed as

$$\epsilon_{long}^{true}(y, t) = \alpha t \left[k(y - y_c)^2 + \epsilon_{long}^{deep}\right] \tag{14}$$

$$\epsilon_{trans}^{true}(y, t) = \alpha t \left[-\nu \left[k(y - y_c)^2 + \epsilon_{long}^{deep}\right]\right] \tag{15}$$

$$\epsilon_{shear}^{true}(x, y, t) = \alpha t \left[kx(y - y_c)\right] \tag{16}$$

$$u_x^{true}(x, y, t) = \alpha t \left[kx(y - y_c)^2 + \epsilon_{long}^{deep} x\right] \tag{17}$$

$$u_y^{true}(y, t) = \alpha t \left[-\nu \left[\frac{k(y - y_c)^3}{3} + \epsilon_{long}^{deep}(y - y_c)\right]\right] \tag{18}$$

The strain field and displacement field were only defined over the region of the image containing the tendon (Figure S3b.). Regions containing tendon were defined semi-automatically with ImageJ (U.S. National Institutes of Health, Bethesda, Maryland, USA) [32].

#### S3.2 Prescribed image deformation and noise addition

Images were artificially deformed by the prescribed displacement field (Equation (17) and (18)) over the region of the tendon using `MATLAB`'s `imwarp` function with linear interpolation settings (Figure S3b.).

The level of noise added to the image was defined by analyzing a series of frames with a stationary participant's arm. The differences between the image intensities of the frames were analyzed, and the mean and standard deviation of the intensity differences were calculated. Across all frames analyzed ($n$ = 2000), it was found that the noise levels remain approximately normally distributed with a mean of 0 and a standard deviation of 10. Therefore, `MATLAB`'s `randn` function was used to add a random distribution of noisy pixel intensities to the frames of the synthetic test case. To match the noise levels measured experimentally, $\mu$ = 0 and $\sigma$ = 10 were set in `randn` (Figure S3c.).

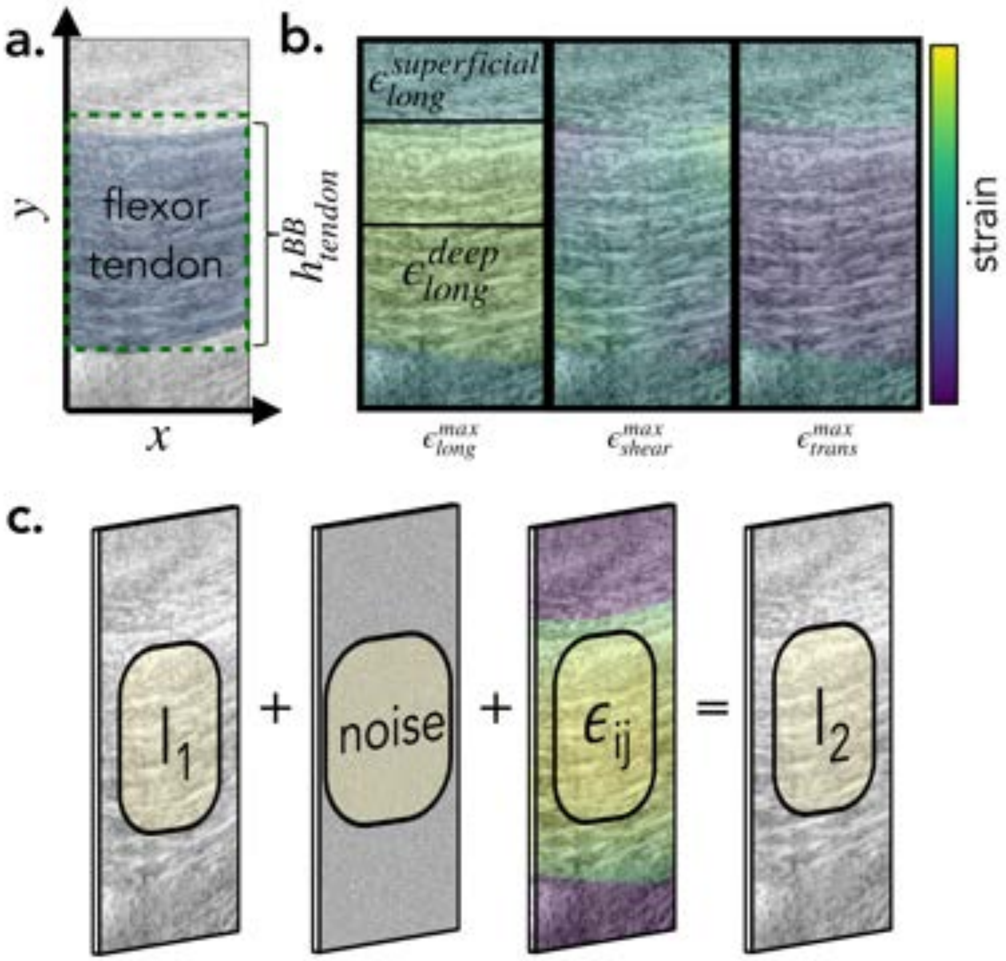

**Figure S3: Diagram of tendon deformation model and ultrasound image processing workflow. a.** A representative ultrasound image containing a flexor tendon, shaded in blue, with a height, $h^{BB}_{tendon}$, green dashed box used in Equation 1. **b.** A strain distribution resulting from the generalized mathematical model, $\epsilon^{max}_{long}$, $\epsilon^{max}_{trans}$, and $\epsilon^{max}_{shear}$, values defined by Equation 9, Equation 10, and Equation 11 respectively, illustrating the regions where these equations were applied. **c.** The general workflow of the image processing pipeline. A reference tendon image, $I_1$, was warped by the generalized mathematical model, noise was introduced, resulting in a deformed image, $I_2$.

### S4 Training set generation

#### S4.1 Overview

To train `StrainNet`, we created a training set containing ultrasound images of tendon deformation with corresponding strain fields. The training set was composed of both synthetically generated images and experimental data, which were processed and combined to ensure a diverse and representative dataset for learning the strain measurement task. Here, we describe the process of generating the training set, including the acquisition of experimental data, image preprocessing, and the combination of synthetic and experimental cases.

#### S4.2 Experimental data acquisition

We conducted an additional, independent ultrasound imaging session using the same protocol described in the experimental protocol. In short, the participant performed a series of contractions, holding each contraction for 5 seconds, followed by a 3-second relaxation period. This protocol allowed us to capture a range of tendon strains during both contraction and relaxation phases. The captured ultrasound images were then used to generate a dataset for the training set creation.

#### S4.3 Image preprocessing

Before incorporating the experimental data into the training set, we performed several preprocessing steps to ensure the images were suitable for training. The preprocessing steps included:

- **Region of Interest (ROI) selection:** The *flexor digitorum superficialis* (FDS) tendon was manually identified and segmented in each ultrasound image. This step focused the training on the relevant tendon deformation while ignoring motion and strain in the surrounding soft tissue.
- **Image normalization:** The pixel intensities of each image were normalized to have a mean of zero and a standard deviation of one. This normalization step facilitated the training process by ensuring consistent image intensity values across the dataset.
- **Augmentation:** To increase the diversity and size of the dataset, we applied various augmentation techniques, including rotations and translations. These augmentations helped the neural network learn to recognize tendon deformation patterns in various contexts and improved the generalization capability of the trained model.

#### S4.4 Application of the generalized mathematical model

The workflow described in Section S3 was combined with the preprocessed experimental data to create a comprehensive training set for `StrainNet`. Specifically, the mathematical model described in Equations (14) - (18) was employed, where the mechanics properties were selected at random. Noise was added to each of the images to emulate real image artifacts, as detailed in Section S3.2. The synthetic cases provided known strain fields, enabling the neural network to learn the relationships between the ultrasound images and the corresponding strain fields. Meanwhile, the experimental data ensured that the neural networks were exposed to real-world tendon deformation patterns.

In total, the training set consisted of 5,000 ultrasound image pairs with known strain fields and added noise—1,250 examples of tensile deformation, compressive deformation, and rigid-body motions for `TensionNet`, `CompressionNet`, and `RigidNet`, respectively (Table S1). All 3,750 examples were used to train the `DeformationClassifier`. Additionally, 1,250 experimental images were included, where the tendon was assumed to undergo tensile deformation, rigid-body motions, and compressive deformation during the contraction, the hold period, and the relaxation, respectively (Table S1). The strain fields corresponding to each image were included into the training set, serving as the ground truth during the training process.

Table S1: Randomly selected mechanical properties for training set generation.

| model | $n$ | $\epsilon_{long}^{sup}$ & $\epsilon_{long}^{deep}$ | $\alpha t$ | $\nu$ | $h_{tendon}^{BB}$ & $y_c$ | exp. aug.$^{\gamma}$ | noise ($\mu \pm \sigma$) |
|---|---|---|---|---|---|---|---|
| `TensionNet` | 1,250 | [2,20] | $[\frac{1}{15}, \frac{1}{5}]$ | $[0.25, 1.5]$ | measured from image$^{\beta}$ | N | $0 \pm 10$ |
| `RigidNet` | 1,250 | [-20,20]$^{\theta}$ | $[\frac{1}{15}, \frac{1}{5}]$ | $[0.25, 1.5]$ | measured from image$^{\beta}$ | N | $0 \pm 10$ |
| `CompressionNet` | 1,250 | [-20,-2] | $[\frac{1}{15}, \frac{1}{5}]$ | $[0.25, 1.5]$ | measured from image$^{\beta}$ | N | $0 \pm 10$ |
| `Deformation Classifier` | 3,750 +1,250$^{\gamma}$ | —pool examples from above— | | | | Y | $0 \pm 10$ |

$^{\theta}$While the images were warped by these largest parameters, the warped image was used for the reference and deformed image.

$^{\beta}$Figure S3a. highlights the measurement of $h_{tendon}^{BB}$ & $y_c$.

$^{\gamma}$ Experimental augmentation whereby experimental images with tensile deformation (during contraction), compressive deformation (during relaxation), and rigid-body motions (during static periods) are included in the training of the `DeformationClassifier`.

#### S5 Synthetic test cases

To benchmark and tune the strain measurement techniques, a series of idealized test cases were created where the applied strain was known. To mimic the experimental procedure detailed in the experimental protocol, the mathematical model outlined in Section S3 was modified to reach its peak after a contraction period of three seconds. This peak was then maintained for five seconds, followed by a relaxation period of three seconds (Figure S4). As a result, the true time-dependent strain field was defined as

$$\epsilon_{long}^{true}(y,t) = \begin{cases} (\frac{t}{3})\epsilon_{long}^{max}(y) & \text{for } 0 \text{ sec} \leq t \leq 3 \text{ sec} \\ \epsilon_{long}^{max}(y) & \text{for } 3 \text{ sec} < t \leq 8 \text{ sec} \\ (1-\frac{t-8}{3})\epsilon_{long}^{max}(y) & \text{for } 8 \text{ sec} < t \leq 11 \text{ sec} \end{cases} \tag{19}$$

$$\epsilon_{trans}^{true}(y,t) = \begin{cases} (\frac{t}{3})\epsilon_{trans}^{max}(y) & \text{for } 0 \text{ sec} \leq t \leq 3 \text{ sec} \\ \epsilon_{trans}^{max}(y) & \text{for } 3 \text{ sec} < t \leq 8 \text{ sec} \\ (1-\frac{t-8}{3})\epsilon_{trans}^{max}(y) & \text{for } 8 \text{ sec} < t \leq 11 \text{ sec} \end{cases} \tag{20}$$

$$\epsilon_{shear}^{true}(x,y,t) = \begin{cases} (\frac{t}{3})\epsilon_{shear}^{max}(y) & \text{for } 0 \text{ sec} \leq t \leq 3 \text{ sec} \\ \epsilon_{shear}^{max}(x,y) & \text{for } 3 \text{ sec} < t \leq 8 \text{ sec} \\ (1-\frac{t-8}{3})\epsilon_{shear}^{max}(x,y) & \text{for } 8 \text{ sec} < t \leq 11 \text{ sec} \end{cases} \tag{21}$$

and the full time-dependent prescribed displacement field was

$$u_x^{true}(x,y,t) = \begin{cases} (\frac{t}{3})u_x^{max}(x,y) & \text{for } 0 \text{ sec} \leq t \leq 3 \text{ sec} \\ u_x^{max}(x,y) & \text{for } 3 \text{ sec} < t \leq 8 \text{ sec} \\ (1-\frac{t-8}{3})u_x^{max}(x,y) & \text{for } 8 \text{ sec} < t \leq 11 \text{ sec} \end{cases} \tag{22}$$

$$u_y^{true}(y,t) = \begin{cases} (\frac{t}{3})u_y^{max}(y) & \text{for } 0 \text{ sec} \leq t \leq 3 \text{ sec} \\ u_y^{max}(y) & \text{for } 3 \text{ sec} < t \leq 8 \text{ sec} \\ (1-\frac{t-8}{3})u_y^{max}(y) & \text{for } 8 \text{ sec} < t \leq 11 \text{ sec} \end{cases} \tag{23}$$

Additionally, studies have shown that tendons exhibit longitudinal strain magnitudes between 4% and 14% under contraction, and therefore the maximum applied strain, which occurs where $y = y_c$, was varied between 4% and 16% [28–31] over five different synthetic test cases (Table S2). We assumed that the tendon was incompressible; therefore, the Poisson's ratio was 0.5 throughout the tendon (Table S2) [33].

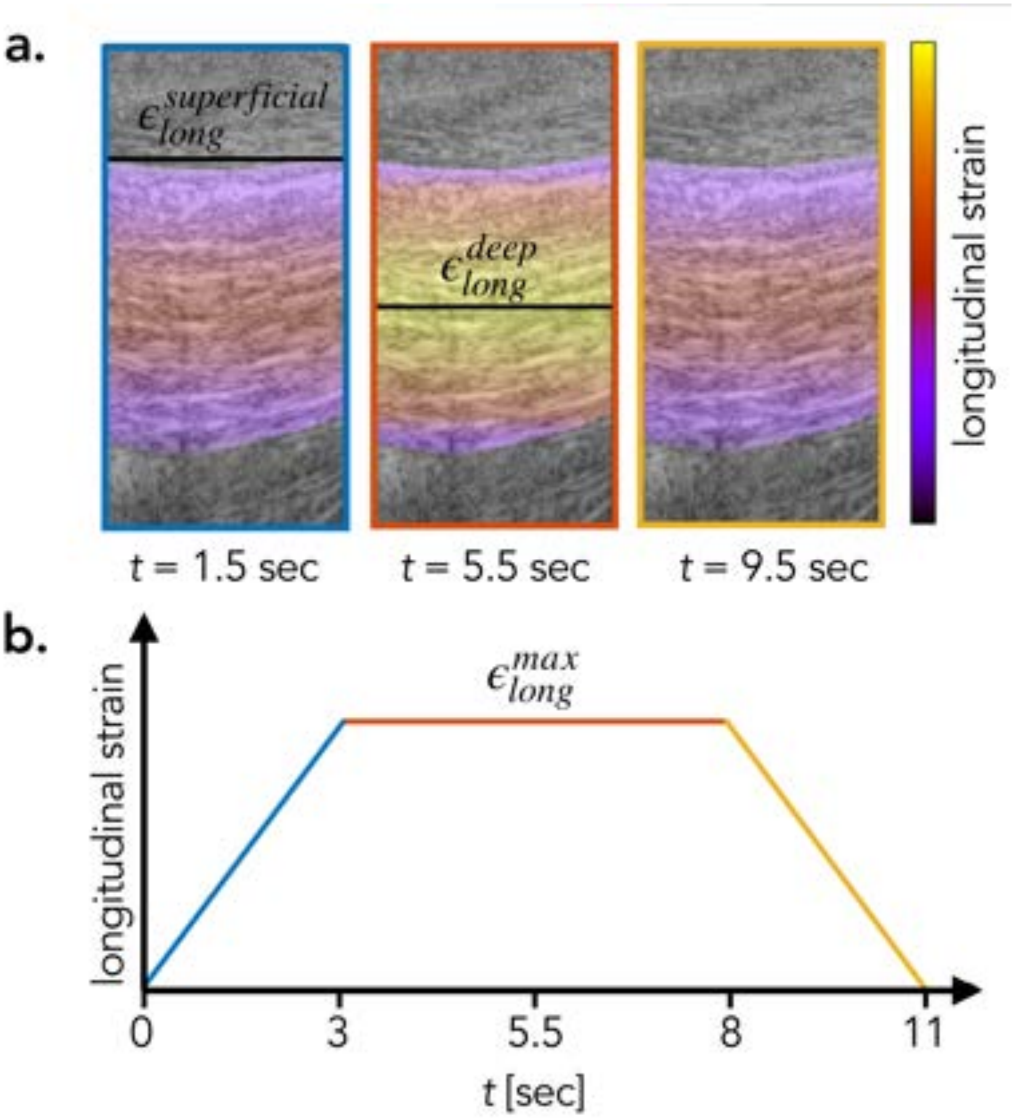

**Figure S4: An overview of synthetic test cases for assessing tendon deformation using ultrasound imaging. a.** Longitudinal strain distribution, $\epsilon_{long}^{true}$, at $t$ = 1.5, 5.5, and 9.5 seconds. The maximal strain occurs at deep layer, $\epsilon_{long}^{deep}$, with superficial layer experiencing 75% of peak strain, $\epsilon_{long}^{superficial}$ [28]. **b.** The time-dependent change in the maximum prescribed longitudinal strain, $\epsilon_{long}^{max}$, over the course of the synthetic test cases. Blue, red, and yellow represent periods of contraction, holding, and relaxation, respectively.

**Table S2: Summary of the five synthetic test cases used to evaluate, tune, and validate the three strain measurement techniques: DIC, DDE, `StrainNet`.**

| Test Case | $\epsilon_{long}^{max}$ | $\epsilon_{long}^{superficial}$ | $\nu$ | Contraction time | Hold time | Relaxation time |
|---|---|---|---|---|---|---|
| 1 | 4% | 3% | 0.5 | 3 sec | 5 sec | 3 sec |
| 2 | 7% | 5.25% | 0.5 | 3 sec | 5 sec | 3 sec |
| 3 | 10% | 7.5% | 0.5 | 3 sec | 5 sec | 3 sec |
| 4 | 13% | 9.75% | 0.5 | 3 sec | 5 sec | 3 sec |
| 5 | 16% | 12% | 0.5 | 3 sec | 5 sec | 3 sec |

#### S6 Reported *in vivo* tendon modulus

The tendon apparent modulus has been a key area of interest of biomechanical researchers. Table S3 provides a comparison of the *in vivo* reported apparent moduli of different human tendons under various loading conditions. The patellar tendon, as investigated by Carroll *et al.*, Reeves *et al.*, and Hansen *et al.*, exhibited apparent moduli of 0.9 GPa, 1.3 GPa, and 1.09 GPa, respectively, under conditions of maximal isometric knee extension, with slight variations in the testing approach. The Achilles tendon, studied by Lichtwark & Wilson and Coombes *et al.*, showed a slightly lower apparent modulus with values of 0.87 GPa and 0.76 GPa, respectively, under the conditions of one leg hopping and maximum voluntary isometric plantarflexion contractions. The tibialis anterior tendon, tested by Maganaris & Paul using 50V applied and maximum isometric load, showed the most varied modulus, with values of 0.45 GPa and 1.2 GPa. Finally, the tendon structures in the vastus lateralis muscle, as investigated by Kubo *et al.*, presented the lowest apparent modulus values of 0.29 GPa and 0.43 GPa under the condition of isometric knee extension torque from a relaxed state to maximum voluntary contraction within 5 seconds. The apparent modulus of these tendons, as shown in Table S3, exhibits considerable variation [34–40].

**Table S3: Comparison of *in vivo* reported mechanical properties of different human tendons under different loading conditions.**

| author | tendon | loading condition | apparent modulus (GPa) |
|---|---|---|---|
| Carroll *et al.* | patellar | maximal 10-s ramp isometric knee extension | $0.9 \pm 0.1$ |
| Reeves *et al.* | patellar | maximal isometric knee extension | $1.3 \pm 0.3$ |
| Hansen *et al.* | patellar | maximal 10-s ramp isometric knee extension | $1.09 \pm 0.12$ |
| Lichtwark & Wilson | achilles | one leg hopping | $0.87 \pm 0.2$ |
| Coombes *et al.* | achilles | maximum voluntary isometric plantarflexion contractions | $0.76 \pm 0.4$ |
| Maganaris & Paul | tibialis anterior | 50V applied & maximum isometric load | $0.45 \pm 0.06$ & $1.2 \pm 0.15$ |
| Kubo *et al.* | tendon structures in the vastus lateralis muscle | isometric knee extension torque from zero (relax) to MVC within 5 s | $0.29 \pm 0.03$ & $0.43 \pm 0.04$ |
